# Protecting synapses from amyloid β-associated degeneration by manipulations of Wnt/planar cell polarity signaling

**DOI:** 10.1101/2020.09.09.273011

**Authors:** Bo Feng, Andiara E. Freitas, Runyi Tian, Yeo Rang Lee, Akumbir S. Grewal, Jingyi Wang, Yimin Zou

**Author notes:** Corresponding Author: Yimin Zou.

## Abstract

Synapse loss is an early event in Alzheimer’s disease and is thought to be associated with amyloid pathology and caused by Amyloid β (Aβ) oligomers. Whether and how Aβ oligomers directly target signaling pathways for glutamatergic synapse maintenance is unknown. Glutamatergic synapse development is controlled by the opposing functions of Celsr3 and Vangl2, core components of the Wnt/planar cell polarity (PCP) signaling pathway, functioning directly in the synapses. Celsr3 promotes synapse formation, whereas Vangl2 inhibits synapse formation. Here we show that oligomeric Aβ binds to Celsr3 and assists Vangl2 in disassembling synapses by disrupting the intercellular Celsr3/Frizzled3-Celsr3 complex, essential for PCP signaling. Together with Vangl2, a Wnt receptor, Ryk, is also required for Aβ oligomer-induced synapse loss in a mouse model of Alzheimer’s disease, 5XFAD, where conditional *Ryk* knockout protected synapses and preserved cognitive function. Our study reveals a fine balance of Wnt/PCP signaling components in glutamatergic synapse maintenance and suggests that overproduced Aβ oligomers may lead to excessive synapse loss by tipping this balance. Together with previous reports that an inhibitor of Wnt/Ryk signaling, WIF1, is found reduced in Alzheimer’s disease patients, our results suggest that the imbalance of PCP signaling in these patients may contribute to synapse loss in Alzheimer’s disease and manipulating Wnt/PCP signaling may preserve synapses and cognitive function.

---

The cause(s) of Alzheimer’s disease has not been well understood due to the complexity and heterogeneity of the disease. The extracellular β-amyloid plaques and intracellular neurofibrillary tangles, formed by hyperphosphorylated tau, are the primary neuropathologic hallmarks for Alzheimer’s disease ^1 2^. They have both been proposed to cause Alzheimer’s disease ^3 4 5 6 7^. More recent findings suggest that a strong but complex involvement of the innate immune system may be a key downstream event to either respond to or enhance amyloid pathology or exacerbate tau-associated pathology. The majority of cases of Alzheimer’s disease are late-onset AD (LOAD), with *APOE4* as the strongest risk factor. Increased Aβ seeding and reduced Aβ clearance appear to be related to the risk, suggesting *APOE4* may affect AD risk, at least in part, by regulating amyloid pathology ^8 9 10^. The relationship between β−amyloid pathology and tauopathy, which corelates better with the progression of cognitive impairment, has not been sorted out as to which one is causal or whether they actually synergize ^11^. Therefore, we elected to focus on an amyloid-dependent event that occurs before tauopathy is fully developed, the initial loss of glutamatergic synapses ^12^.

Amyloid precursor protein and its metabolites regulate synaptic transmission, plasticity and calcium homeostasis ^13^. While picomolar amounts of Amyloid β (Aβ) is essential for long-term potentiation (LTP) and long-term depression (LTD), nanomolar concentrations of Aβ inhibits LTP induction. Therefore, Aβ is thought to be a negative feedback mechanism to reduce neural activity in neuronal networks. Aβ readily self-associates to form a range of neurotoxic soluble oligomers and insoluble deposited fibers ^14^. Soluble Aβ oligomers induce loss of glutamatergic synapses, loss of LTP and decrease of dendritic spine density ^15 16 17 18 19 20^. Although several receptors have been found to bind to Aβ oligomers regulating synaptic plasticity, including cellular prion protein (PrP^C^), EphB2 and paired immunoglobulin-like receptor B (PirB) (or its human ortholog leukocyte immunoglobulin-like receptor B2 (LilrB2)) ^21–23^, how Aβ oligomers directly mediates glutamatergic synapse loss is not known.

Planar cell polarity (PCP) signaling components play essential roles in glutamatergic synapse formation in development ^24^. Frizzled3 is enriched in the synaptic vesicles and on the plasma membrane of the presynaptic boutons and Vangl2 is enriched on the plasma membrane of the postsynaptic neurons and in the postsynaptic density (PSD), whereas Celsr3 is on the plasma membranes of both pre- and postsynaptic neurons. Celsr3 promotes synapse formation, whereas Vangl2 inhibits synapse formation ^24^. Whether PCP signaling plays a role in mature neural circuits in adulthood is not known. We show here that the PCP components and a Wnt receptor, Ryk, are also present in the adult glutamatergic synapses and that Aβ oligomers are unable to cause synapse loss when *Vangl2* is conditionally knocked out *in vitro* and *in vivo* in adult hippocampus. Vangl2 disassembles glutamatergic synapses by disrupting the intercellular complex formed by Celsr3/Frizzled3-Celsr3, essential for PCP signaling. Aβ oligomers bind to Celsr3 and assists Vangl2 in disrupting this intercellular complex. Ryk regulates PCP signaling by directly interacting with and promoting the function of Vangl2 ^25 26^. A function of Ryk in regulating the stability of neuronal synapses has not been reported. We found that Ryk is also required for Aβ oligomer-induced synapse loss *in vitro* and *in vivo* as shown by a function-blocking monoclonal Ryk antibody and *Ryk* conditional knockout. Conditional *Ryk* knockout prevented loss of synapses and preserved cognitive functions of the *5XFAD* mice. In Alzheimer’s disease patients, an inhibitor of Wnt/Ryk signaling, WIF1, is found greatly reduced, suggesting that the imbalance of the Wnt/PCP signaling pathway may contribute to synapse loss in Alzheimer’s disease and the Wnt/PCP pathway may be a novel therapeutic target ^27 28^.

## RESULTS

### Wnt/PCP signaling is essential for synapse maintenance and synapse number control in adulthood and may be a target for synapse degeneration *in vivo*

PCP components are essential regulators of glutamatergic synapse formation during early postnatal development ^24^. To test whether they also play a role in the mature nervous system, we first examined their expression and localization in adult hippocampus (8 weeks) (Fig. 1a). Like observed in postnatal day 14, we found that, Celsr3 and Vangl2 are also colocalized in the glutamatergic synaptic puncta as visualized by costaining with Bassoon and PSD95 in the hippocampus at 2 months of age. Whether the Wnt receptor, Ryk, is also localized in glutamatergic synapses is not known. We found that Ryk protein is present in adult synapses (Fig. 1a). Therefore, our new results indicate that Wnt-Ryk/PCP components maintain their expression in the adult glutamatergic synapses (Fig. 1b). To assess whether PCP signaling continues to regulate synapse maintenance in adulthood, we used the CRISPR-cas9 system to knock out all three *Celsrs* to avoid compensation. We designed single-guide RNAs (sgRNAs) targeting *Ceslr1, Celsr2 and Celsr3*. CRISPR-induced genome editing was verified in Neuro2A cells (Extended data Fig.1a) and knockout efficiency was verified with Western blots of protein extracts from cultured hippocampal neurons (Extended data Fig.1b). One month after injecting the *AAV sgRNA* into the CA1 region of the *Cas9* mice, we analyzed the synapse numbers by costaining with synaptic makers and found that *AAV-sgCelsr1,2,3* significantly reduced the synapse numbers in the stratum radiatum of the adult hippocampus (Extended data Fig.1c,d). Therefore, *Celsrs* are essential for synapse maintenance in adulthood.

**Fig. 1.**
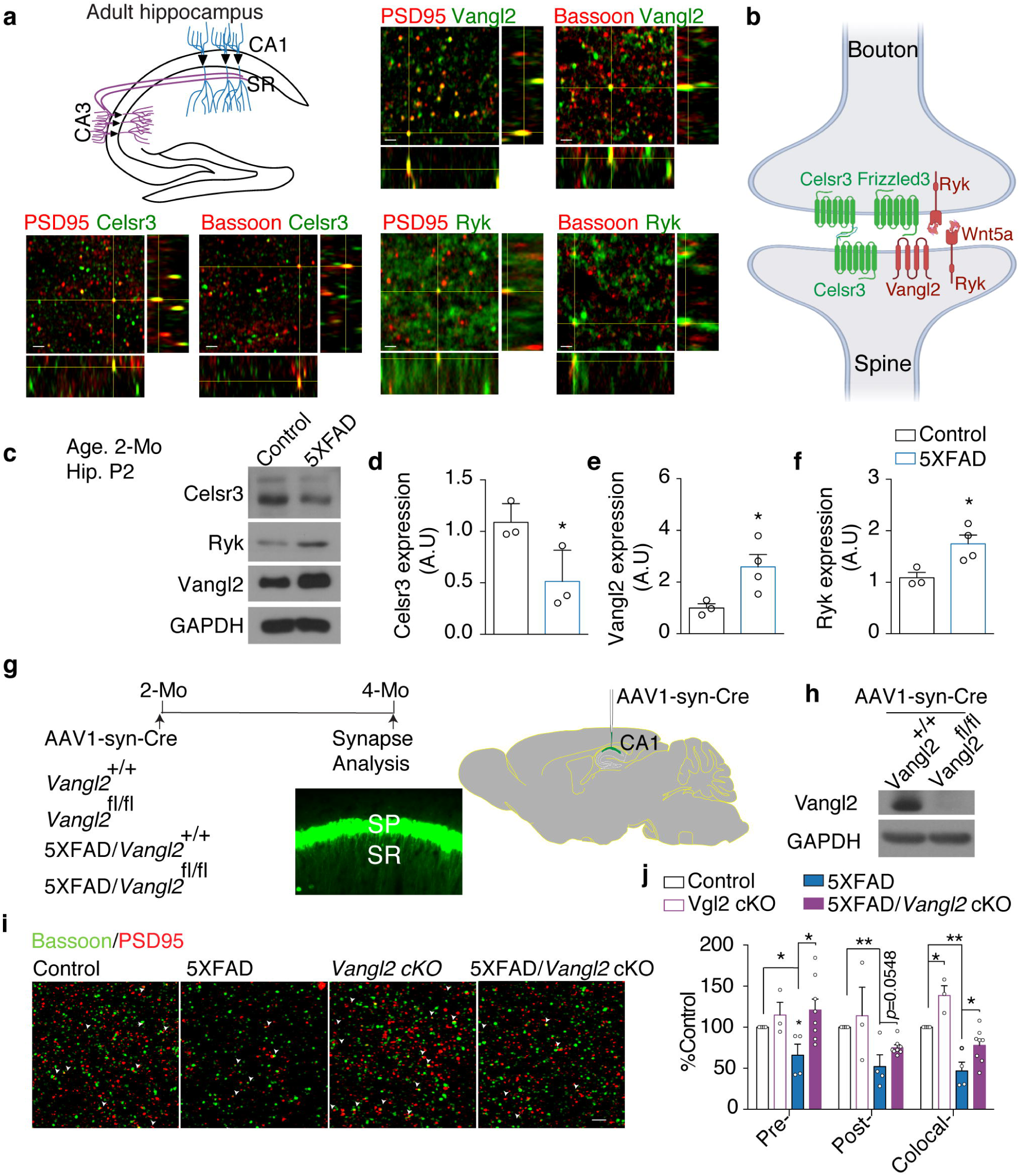
Localization of Wnt/PCP signaling components in glutamatergic synapses in adult hippocampus and requirement of *Vangl2* for synapse loss in the *5XFAD* mice. **a**, Costaining of Vangl2 (Green) and Celsr3 (Green) with postsynaptic markers, PSD-95 (Red) and presynaptic mater bassoon (Red). Arrows indicate colocalized puncta. SR, stratum radiatum; SP, stratum pyramidale. **b,** Schematic diagram showing the distribution of the Wnt/PCP signaling components in glutamatergic synapses. **c-f,** Expression levels of Celsr3, Ryk and Vangl2 in the P2 synaptosome fraction for adult hippocampus in control and 5XFAD transgenic mice. Student *t*-test. **g,** Schematics illustrating the experimental design. *AAV1-hSyn-eGFP-Cre* virus was injected into CA1 region of the hippocampus bilaterally. 2 months later, animals were fixed with perfusion and sectioning and stained with synaptic markers. **h,** Vangl2 protein level in total protein extracts from hippocampi injected with *AAV1-hSyn-eGFP-Cre* virus. **i,** Representative images of costaining for Bassoon (red)- and PSD95 (green)- (arrowheads indicate colocalization) in the stratum radiatum. **j,** quantification of (I). n=5 for control mice, n=3 for *Vangl2 cKO* mice, n=4 for *5XFAD* mice and n=8 for *5XFAD*; *Vangl2 cKO* mice. One-way ANOVA. \**P* < 0.05, ** *P* < 0.01. Scale bar 1 μm in (**a**), scale bar: 2 μm in (**i**). Error bars represent SEM.

We then tested whether the Wnt/PCP signaling is altered in models *in vivo* synapse loss. Synapse loss is well documented in the *5XFAD* transgenic mice ^29^. We found that, indeed, Celsr3 protein level was significantly reduced, whereas the levels of Vangl2 and Ryk were increased in the P2 synaptosome fraction extracted from the hippocampi of the adult *5XFAD* transgenic mice (Fig. 1c-f). Because Vangl2 inhibits synapse formation ^24^, we crossed *5XFAD* with the *Vangl2* conditional knockout (*cKO*) line and injected *AAV1-hSyn-eGFP-Cre* (Adeno-associated virus (AAV) that harbors the human synapsin (*hSyn*) promotor with cytomegalovirus (*CMV*) enhancer driving the Cre recombinase) to the CA1 region of the adult hippocampus at the age of 2 months and then analyzed the synapse numbers at the age of 4 months (Fig. 1g,h). We observed an 27% increase of synapse numbers in *Vangl2 cKO* (first two columns in colocalized), suggesting that *Vangl2* negatively regulate synapse numbers in adulthood *in vivo*. We found that the synapse number was 66% higher in *5XFAD* crossed with Vangl2 cKO (magenta column in colocalized) than *5XFAD* alone (blue column in colocalized) (Fig. 1i,j). Therefore, *Vangl2* cKO rescued 39% of the synapses. The synapse number analyses were done in double blind.

### *Vangl2* is required for Aβ oligomer-induced synapse loss *in vitro* and *in vivo*

Because *5XFAD* is a model for amyloid-β overexpression, we first tested whether *Vangl2* is required for amyloid-β oligomer (Aβ oligomer)-induced synapse loss in cultured hippocampal neurons isolated from embryonic day 18 (E18.5) of *Vangl2 cKO* mice. *AAV1- hSyn-eGFP-Cre* was added into the culture on the 7^th^ day after the start of culture (DIV-7). At DIV14, 400 nM monomer equivalent of Aβ oligomers were added and the cultures were fixed 12 hours later and stained and analyzed (Fig. 2a). Vangl2 protein level was found to be significantly reduced in *Vangl2*^fl/fl^ infected with the *AAV1-hSyn-eGFP-Cre* (Fig. 2b). Extended data Fig.2 shows that the effective concentration of dimer added was 80 nM and tetramer was ∼152 nM. The calculation of concentration is described in the Methods ^21, 30^. Littermate control neurons (*Vangl2^+/+^*, Control) were also treated with *AAV1-hSyn-eGFP-Cre*. We found that, in control neurons, Aβ oligomers reduced synapse numbers by 30% as shown by costaining for Bassoon and PSD-95 (Fig. 2c,d). However, the number of glutamatergic synapses in *Vangl2* cKO neuron was unchanged compared with *Vangl2* cKO neurons not treated with Aβ oligomers (Fig. 2c,d). Consistent with our previous finding that Vangl2 inhibits synapse formation, *Vangl2 cKO* itself lead to 40% increase of synapse numbers during the 7.5 days of culture of the E18.5 embryonic neurons (Fig. 2c,d) ^24^.

**Fig. 2.**
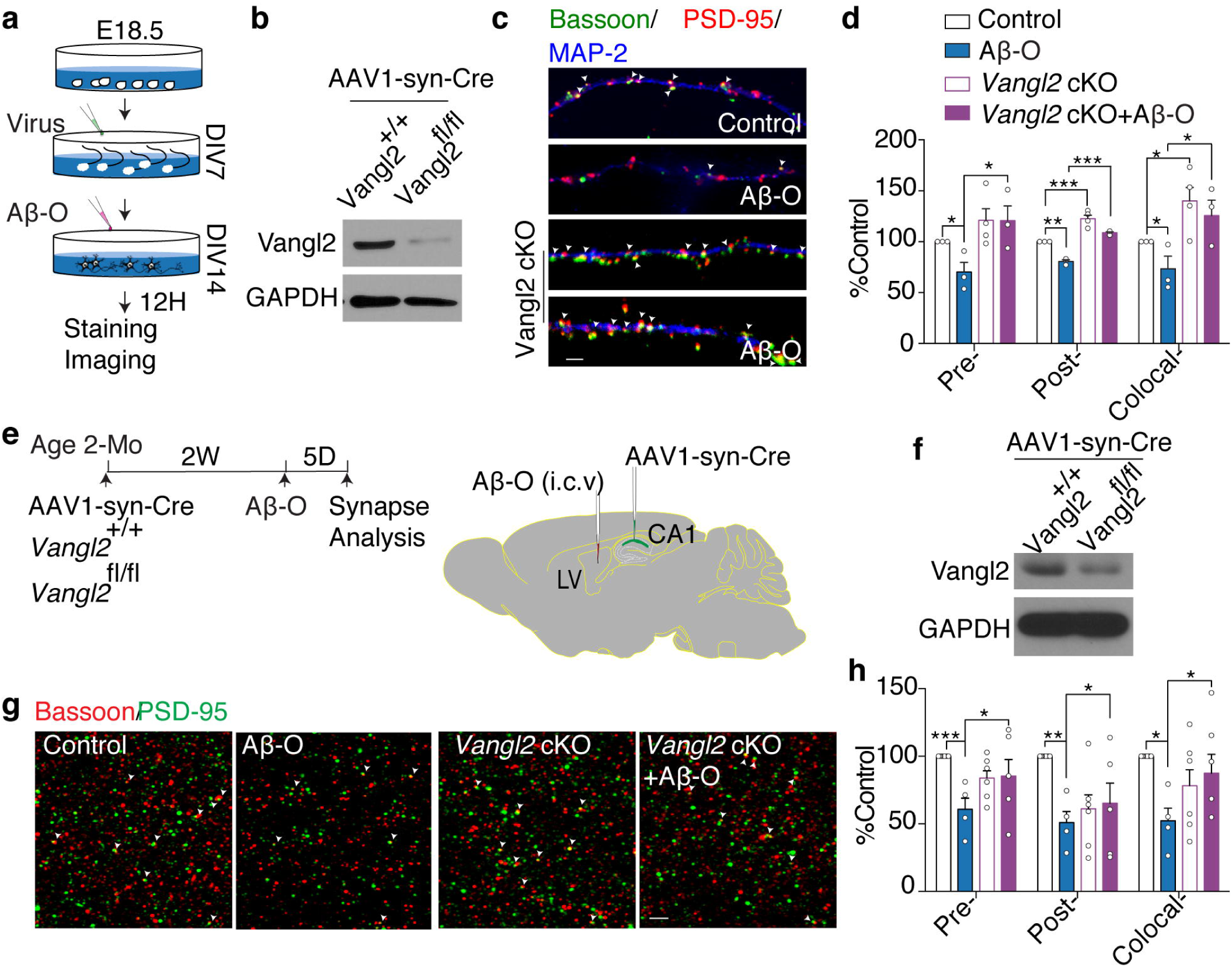
Vangl2 is required for Aβ oligomer-induced synapse loss *in vitro* and *in vivo*. **a,** Schematics illustrating the experimental design. *AAV1-hSyn-eGFP-Cre* virus was added to hippocampal neuron cultures on DIV7 for 7 days and then oligomeric Aβ42 was added. 12 hours later after adding oligomeric Aβ42, cultures were fixed for staining synaptic markers. **b,** Western blot showing the level of Celsr3 and Vangl2 proteins in cultures infected with the *AAV1-hSyn-eGFP-Cre* virus. **c,** Immunostaining for pre-(green) and postsynaptic (red) puncta of glutamatergic synapses (arrowheads) in 14-DIV hippocampal cultures from littermate *Vangl2*^+/+^ or *Vangl2*^fl/fl^ with or without oligomeric Aβ42. **d,** Quantification of (**c**). n=3 for *Vangl2*^+/+^ mice, n=4 for *Vangl2*^fl/fl^ from 3 independent experiments. **e,** Schematics illustrating the experimental design. *AAV1-hSyn-eGFP-Cre* virus was injected into CA1 region of the hippocampus bilaterally. 2 weeks later, oligomeric Aβ was injected into cerebroventricular. 5 days after Aβ oligomer injection, animals were fixed with perfusion and sectioning and stained with synaptic markers. **f,** Vangl2 protein level in the total hippocampus extract from animals injected with the *AAV1- hSyn-eGFP-Cre* virus. **g,** Representative images of Bassoon (red)- and PSD95 (green)- immunoreactive puncta (arrowheads) in stratum radiatum of *Vangl2*^+/+^ and *Vangl2*^fl/fl^ hippocampus (CA1) with or without oligomeric Aβ injection and quantification of synapse numbers. **h,** Quantification of (**g**). One-way ANOVA. n=8 of *Vangl2*^+/+^ mice, n=3 of *Vangl2*^+/+^ mice with oligomeric Aβ injection, n=6 of *Vangl2*^fl/fl^ mice and n=5 of *Vangl2*^fl/fl^ mice with oligomeric Aβ injection. \**P* < 0.05, \*\**P* < 0.01, One-way ANOVA. Scale bar: 2 μm in (**c**) and (**g**). Error bars represent SEM.

To test whether *Vangl2* is required for Aβ oligomers-induced synapse loss *in vivo* and in adulthood, we first injected *AAV1-hSyn-eGFP-Cre* into the hippocampal CA1 region of *Vangl2*^+/+^ (Control) and *Vangl2*^fl/fl^ mice at the age of 2 months. 2 weeks later, we injected 5 ng of Aβ oligomers into cerebral ventricles bilaterally ^31^ (Fig. 2e). 5 days after injection of Aβ oligomers, animals were perfused and the synapse number was analyzed in blind. Vangl2 protein level was found significantly reduced in *Vangl2 cKO* (Fig. 2f). We anticipated that the extent of rescue from acute intracerebroventricular injection of Aβ oligomers in *Vangl2 cKO* animals examined 5 days after the injection of Aβ oligomers may be more than conditionally knocking out of *Vangl2* in *5XFAD* transgenic mice (Fig. 1i,j). In the *5XFAD* experiment, Aβ oligomers and other pathological factors were already present before *Vangl2* was conditionally knocked at the age of two months and may have caused synapse loss or changes of synaptic structures. Indeed, we observed a significant loss of synapses in control animals injected with Aβ oligomers but not in the *Vangl2 cKO* mice injected with Aβ oligomers and *Vangl2* preserved the synapse number to the extent that is comparable to the control animals (Fig. 2g,h). In contrast to the experiment with cultured neurons from E18.5 embryos, *Vangl2* cKO in adulthood itself showed no significant changes in synapse numbers 19 days weeks after the *AAV1-hSyn-eGFP-Cre* injection (Fig. 2g,h). It might because that synapse turnover is not as rapid in adulthood such that 19 days of time is not long enough to observe synapse number changes. However, 2 months would be long enough time to observe an increase of synapse numbers in *Vangl2 cKO* (Fig. 1i,j).

### Binding of Aβ oligomers to Celsr3 is required for Aβ oligomer-induced synapse loss

To understand how PCP signaling may mediate Aβ oligomer-induced synapse loss, we investigated a potential direct interaction between Aβ oligomers and the PCP components. PCP components are localized in glutamatergic synapses in similar fashions as in the asymmetric epithelial cell-cell junctions during PCP signaling (Fig. 1a,b) ^24^. Among the 6 core PCP components, Celsrs, Frizzleds and Vangls are present on the plasma membrane. To determine whether Aβ oligomers target any one(s) of those proteins, we measured binding of biotin-Aβ42 oligomers to HEK293T cells expressing mouse Vangl2 (Vangl2- Flag), Frizzled3 (Frizzled3-HA), Celsr3 (Celsr3-Flag) or control vector (pCAGEN). We found that Aβ oligomers only bound to Celsr3, but not Vangl2 or Frizzled3 (Fig. 3a), with an apparent dissociation constant (*K*d) of ∼40 nM equivalent of total Aβ peptide (Fig. 3b). Aβ monomer did not bind to Celsr3 (Extended data Fig. 3).

**Fig. 3.**
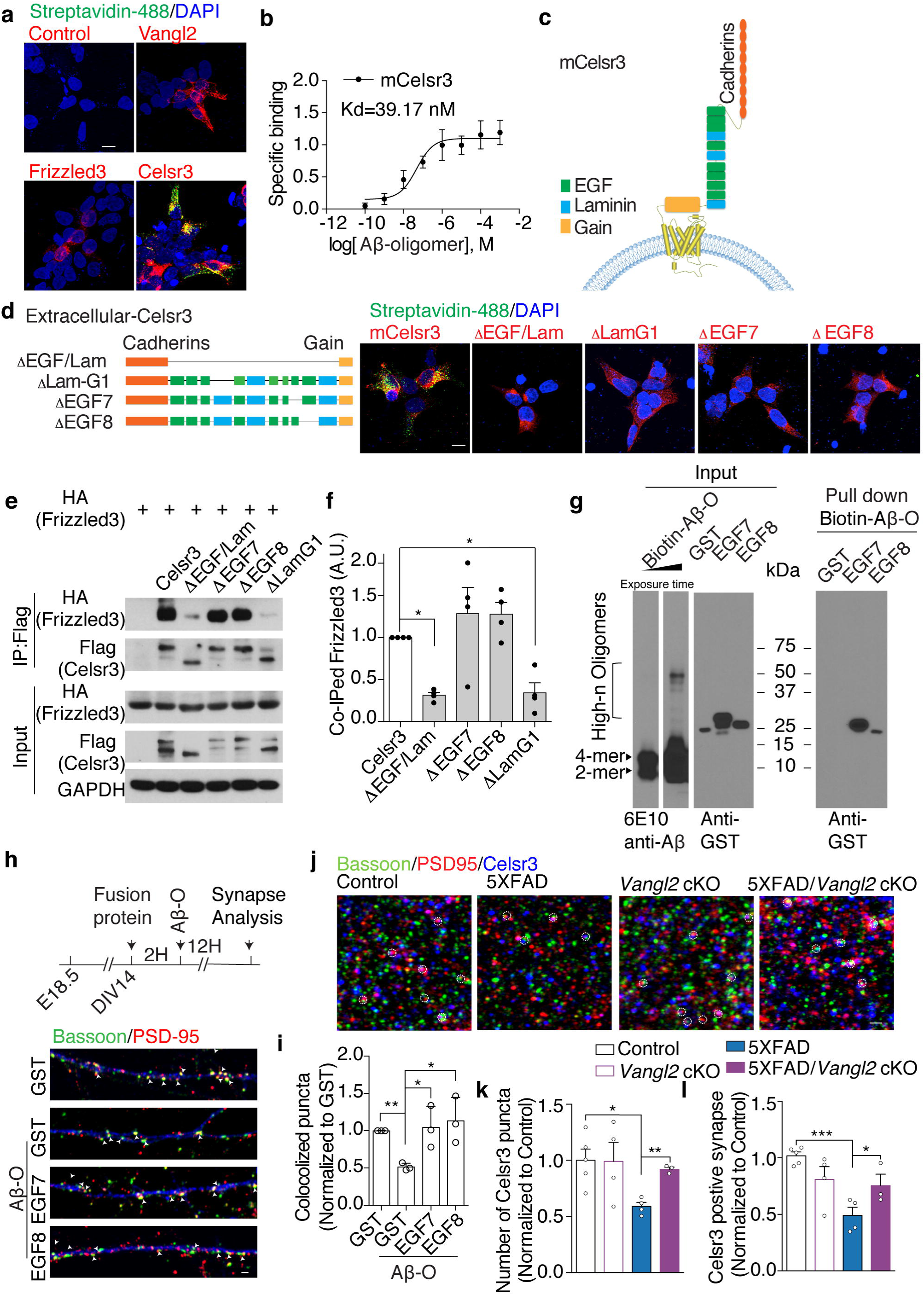
Celsr3 is the binding protein for oligomeric Aβ and the binding of oligomeric Aβ to the EGF7 and EGF8 domains of Celsr3 is required for synapse loss. **a**, Staining of the HEK293T cells transfected with Vangl2-Flag, Frizzled3-HA, Celsr3- Flag, or control vector (pCAGEN) and incubated with oligomeric Aβ42 (200 nM total peptide, monomer equivalent). Bound oligomeric Aβ42 (green) was visualized using 488- conjugated streptavidin. **b,** Binding curve of Oligomeric Aβ42 to Celsr3-expressing HEK293T cells (concentration showed as total peptide, monomer equivalent). **c,** Schematics of mouse Celsr3 extracellular domain with 9 cadherin domains, 8 EGF domains and 3 laminin domains. **d,** Staining of bound Aβ42 (200 nM total peptide, monomer equivalent) with Celsr3-Flag-transfected or truncated Celsr3-Flag transfected HEK293T cells. Bound oligomeric Aβ42 (green) was visualized using 488-conjugated streptavidin. Scale bar 10 μm. **e-f,** Immunoprecipitated assays testing the interaction between Frizzled3 and Celsr3 or with truncated Celsr3 and quantification data of the expression level of co-IPed Frizzled3. \**P* < 0.05. One-way ANOVA. Means ± SEM. **g,** Pulldown assay for purified GST function proteins mixed with biotinylated Aβ42 oligomers by Streptavidin. **h- i,** Aβ oligomer-induced synapse loss in the presence of EGF7-GST or EGF8-GST fusion proteins. **J-l,** Staining and quantification of Celsr3 positive puncta and glutamatergic synapses (Celsr3 colocalized with PSD95 and bassoon). \**P* < 0.05, ** *P* < 0.01. Student *t*- test. Scale bar: 10 μm in (**a**) and (**d**). Scale bar: 1 μm in (**h**) and (**j**). Error bars represent SEM.

Celsr3 belongs to the family of adhesion G-protein coupled receptors (GPCRs) with a large extracellular region, which contains 9 cadherin domains, 8 EGF repeats and 3 laminin domains (Fig. 3c). Cadherin domains are for homophilic binding. To determine the domains of Celsr3 responsible for binding of Aβ oligomers, we first made a deletion construct that lacks all the EGF repeats and Laminin domains and tested binding in HEK293T cells. We found that Aβ oligomers did not bind to this truncated protein, suggesting that Aβ oligomers do not bind to the cadherin domains but rather bind to the EGF repeats and the Laminin domains (Fig. 3d and Extended data Fig. 4a). We then made a series of Celsr3 constructs that lack the individual EGF and laminin domains (Extended data Fig. 4b) and tested for binding to Aβ oligomers in HEK293T cells. We found that two EGF domains, EGF7 and EGF8 and one Laminin domain, Lamnin-G1 are required for binding of Aβ oligomers (Fig. 3d and Extended data Fig. 4c). The human homolog of murine Celsr3 also contains 9 cadherin domains, 8 EGF repeats and 3 laminin domains.

The Laminin G1 and EGF7 domains of *h*Celsr3 aligns closely with that of *m*Celsr3 with homology of 98.537% and 80%, respectively. The amino acid sequence of the EGF8 domain of *h*Celsr3 is 100% homologous with that of the EGF8 domain of *m*Celsr3 (Extended data Fig. 5a). We found that Aβ oligomers also bound to *h*Celsr3 with an apparent dissociation constant (*K*d) of ∼70 nM equivalent of total Aβ peptide (Extended data Fig. 5b). Like with the *m*Celsr3, EGF7 and EGF8 and one Laminin domain, Lamnin- G1 of *h*Celsr3 are required for binding with Aβ oligomers (Extended data Fig. 5b).

In PCP signaling, protein-protein interaction is essential for the establishment of cell and tissue polarity along the tissue plan. Celsr3 forms a complex with Frizzled3 on the plasma membrane of one cell, and Celsr3 forms a complex with Vangl2 on the plasma membrane of the neighboring cells. We then tested whether the Aβ oligomer-binding domains of Celsr3 are involved in the protein-protein interactions among PCP components. We expressed Frizzled3 or Vangl2 together with wild type Celsr3 or mutant Celsr3 (with domain deletions) in HEK293T cells. We found that deletion of all 8 EGF repeats and 3 Laminin domains caused a 68% reduction of the interaction between Celsr3 and Frizzled3. Deletion of Laminin G1 lead to 66% reduction of the interaction between Celsr3 and Frizzled3, whereas deleting EGF7 or EGF8 did not affect the interaction between Celsr3 and Frizzled3 (Fig. 3e,f). The interaction between Vangl2 and Celsr3 did not require EGF repeats or Laminin domains (Extended data Fig. 6). To ask whether the binding of Aβ oligomers on these domains is important for synapse loss, we sought to examine whether these domains can block Aβ oligomer-induced synapse loss. We tested the role of EGF7 and EGF8 domains, which are not required for Frizzled3 interaction. We performed binding assays with purified EGF7-GST or EGF8-GST fusion proteins and found that both domains can bind to Aβ oligomers, as pulling down biotinylated Aβ oligomers with streptavidin (NeutrAvidin agarose) can pull down the EGF7-GST and EGF8-GST fusion proteins (Fig. 3g). We then added EGF7-GST or EGF8-GST fusion proteins to neuronal hippocampal culture and found that they both blocked Aβ oligomer-induced synapse loss (Fig. 3h,i). Therefore, the EGF7 and EGF8 domains are likely the direct target of Aβ oligomers. To further assess whether Celsr3 is the target of Aβ oligomer-induced synapse loss, we tested whether the Celsr3 positive synapses were lost in the 5XFAD transgenic mice and whether they are rescued by *Vangl2* cKO. We found that in in 5XFAD transgenic mice, Celsr3 puncta and Ceslr3 positive synapses were reduced, both of which were rescued by *Vangl2* cKO (Fig. 3j-l).

### Aβ oligomers enhance the function of Vangl2 in disrupting the intercellular complex essential for PCP signaling

To address how the binding of Aβ oligomers with Celsr3 may lead to synapse loss, we performed a series of biochemistry experiments. PCP signaling is known to be mediated by a set of dynamic protein-protein interactions. One such essential interaction is an asymmetric intercellular complex made of Frizzled and Celsr (Fmi) ^32^. The Frizzled/Celsr complex on the plasma membrane on the distal side of a cell forms an intercellular complex with Celsr on the plasma membrane on the proximal side of the neighboring cell (distal to the first cell) bridging the two cells via the homophilic interaction of the cadherin repeats of the two Celsr proteins. Such an asymmetric intercellular bridge was shown to be sufficient to polarize both cells even in the absence of Van Gogh (Vangl) and, therefore, is thought to be essential for PCP signaling ^32^.

To ask whether and how Vangl2 may negatively regulate synapse numbers, we established an assay to test the intercellular PCP complex, similar to the “transcellular interaction assay” ^33^. We co-transfected Frizzled3 (HA-tagged) and Celsr3 (untagged) in one dish of HEK293T cells and transfected Celsr3 (Flag-tagged) and Vangl2 in another. After culturing them separately for one day, we mixed them together and cultured for one more day and then performed co-immunoprecipitation to test protein-protein interactions (Fig. 4a). To test whether Vangl2 disrupts the intercellular bridge, we pulled down Frizzled3 and test how much Celsr3 from the neighboring cell was coimmunoprecipitated. We found that Vangl2 disrupted this intercellular complex as much less Flag-tagged Celsr3 was pulled down by HA-tagged Frizzled3 (Fig. 4b,c). To test the role of Vangl2, we transfected 1ug of the Vangl2 expression construct (Fig. 4b,c). We think Vangl2 is likely to disrupt this intercellular complex by weakening the interaction between Ceslr3 and Frizzled3, because the presence of Vangl2 alone from the neighboring cell caused the reduction of the interaction between Frizzled3 and Celsr3 by 30-40% (Fig. 4d,f) and that Celsr3 from neighboring cell does not affect the complex between Frizzled3 and Celsr3 (Extended data Fig. 7).

**Fig. 4.**
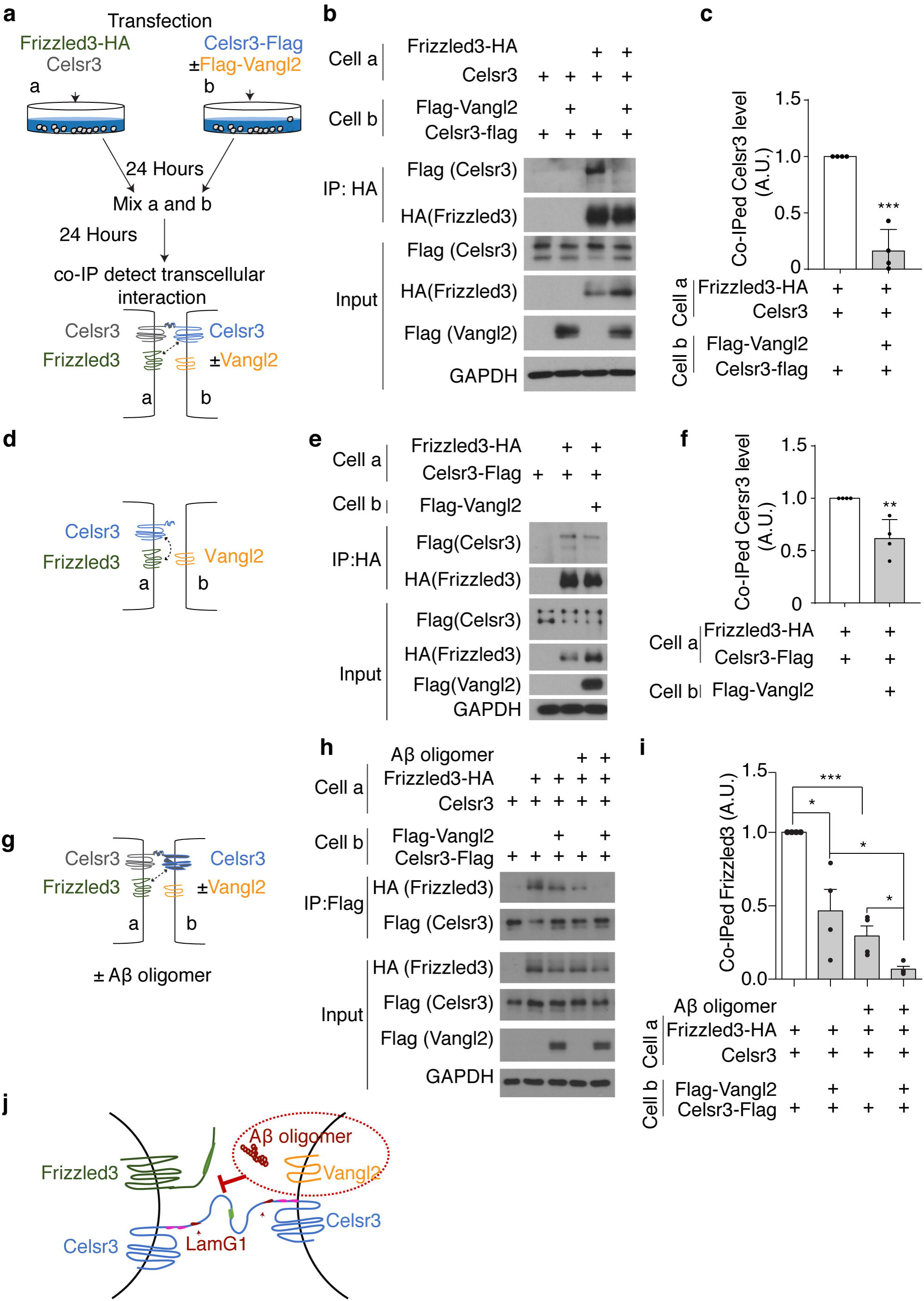
Aβ oligomer enhances Vangl2’s function in disrupting the intercellular complex of Celsr3/Frizzled3-Celsr3. **a**, Schematics illustrating the experimental design of the “transcellular interaction” assay. **b,** Testing the intercellular complex by immunoprecipitating Frizzled3 in one cell and detecting Celsr3 in the neighboring cell with or without Vangl2. **c,** quantification (**b**). **d-f,** The effect of Vangl2 from one cell on the interaction between Frizzled 3 and Celsr3 in the neighboring cells. **g-i,** The effect of Oligomeric Aβ on the function of Vangl2 in disrupting the intercellular complex between Celsr3/Frizzled3 in one cell and Celsr3 in the neighboring cell. **j,** Working hypothesis. Oligomeric Aβ42 assists Vangl2 in weakening the interaction between Celsr3 and Frizzled3 by binding with Celsr3-Laminin G. \**P* < 0.05, \*\**P* < 0.01 and \*\*\**P* < 0.001. One-way ANOVA. Western blot results are representative of four biological replicates. Error bars represent SEM.

In order to determine how Aβ oligomers lead to synapse loss, we tested whether and how Aβ oligomers promote the function of Vangl2 in disrupting the intercellular complex. First, we found Aβ oligomers did not disrupt the interaction between Celsr3 and Frizzled3 that were transfected and expressed in the same cell, suggesting that Aβ oligomers themselves are not sufficient to disrupt the Ceslr3-Frizzled3 complex (Extended data Fig. 8a-c). As shown previously, Vangl2 expressed in neighboring cell can decrease the interaction between Celsr3 and Frizzled 3 by 30-40% (Fig. 4d,f). The interactions between Frizzled3 and Celsr3 were reduced to a greater extent when Aβ oligomers were added to this culture (Extended data Fig. 8d,f), indicating Aβ oligomers may enhance the function of Vangl2 in disrupting the Celsr3-Frizzled3 complex across the cell-cell junction. Furthermore, we found that Aβ oligomers can also disrupt the intercellular complex, as the HA-tagged Frizzled3 in one cell pulled down much less Flag-tagged Celsr3 from a neighboring cell when Aβ oligomers were added to the mixed culture (Extended data Fig. 8g,i). This suggests that the intercellular interaction of Celsr3 between two neighboring cells may weaken the intracellular interaction between Celsr3 and Frizzled3 within the same cell, allowing Aβ oligomers to more efficiently disrupt the entire intracellular interaction. In glutamatergic synapses, Celsr3 is present on both pre- and post-synaptic sides. Finally, we found that adding Aβ oligomers to mixed cultures with Vangl2 expressed with Celsr3 lead to the greatest disruption of this intercellular complex (Fig. 4g-i). In order to test the role of Aβ oligomers in enhancing the function of Vangl2, we transfected 0.5 ug of Vangl2 expression construct so that Vangl2 is at suboptimal concentration (Fig. 4g-i). Therefore, we propose that Aβ oligomers enhance the function of Vangl2 by disrupting the Celsr3/Frizzled3 intracellular complex and thus the asymmetric Celsr3/Frizzled3-Celsr3 intercellular complex essential for PCP signaling. This is probably because the binding of Aβ oligomers to the Laminin G1 domain of Celsr3 weakens the interaction between Celsr3 and Frizzled3, allowing Vangl2 to more efficiently disrupt the asymmetric intercellular complex of Celsr3/Frizzled3-Celsr3 and thus disassemble more synapses (Fig. 4j).

### The Wnt/Ryk/Vangl2 signaling axis mediates Aβ oligomer-induced synapse loss *in vitro* and *in vivo*

In order to further address how Vangl2 mediates Aβ oligomer-induced synapse loss and identify potential therapeutic targets to protect synapses and preserve functions, we explored regulators of core PCP signaling components. As Ryk is a coreceptor for Wnt in PCP signaling by directly interacting with Vangl2 and promoting the function of Vangl2, we sought to test whether Wnt/Ryk signaling is involved in Vangl2 function in this context^25 26^. Wnt5a cause reduction of synapse numbers probably by causing Frizzled3 endocytosis ^34 24^. We first tested whether Ryk mediates Wnt5a function in regulating synapse numbers and whether Ryk does so in a Vangl2-dependent manner. Hippocampal neurons isolated from E18.5 WT mice were treated with Wnt5a on DIV14 for 12 hours or pre-treated with a function blocking monoclonal Ryk antibody, which blocks the binding between Wnts and Ryk, for 2 hours (Fig. 5a) ^35^. Normal mouse IgG was used as control. Wnt5a caused 30% reduction in the number of colocalized puncta. In contrary, Wnt5a did not produce a significant difference in synapse number if the cultures were pre-treated with Ryk antibody (Fig. 5a,b), suggesting that Wnt5a inhibits synapse formation through binding to Ryk as the receptor. During the 14 hours of culture time, Ryk antibody itself did not cause any significant change in synapse numbers (Fig. 5b). To test whether Vangl2 mediates the inhibitory function of Wnts downstream of Ryk, we cultured *Vangl2*^+/+^ (Control) and *Vangl2 cKO* embryonic hippocampal neurons (infected with *AAV1-hSyn-eGFP-Cre*) and treated neurons with Wnt5a on DIV 14 for 12 hours. We found that Wnt5a addition to *Vangl2*^+/+^ caused a 30% reduction in the number of colocalized puncta, whereas Wnt5a addition to *Vangl2 cKO* neurons did not produce a significant difference compared with untreated *Vangl2 cKO* neurons (Fig. 5c,d), suggesting that *Vangl2* is required for the inhibitory function of Wnt5a in synapse formation. Consistent with our previous finding (Fig. 2c,d), *Vangl2* cKO itself lead to more synapses 7.5 days after the introduction of Cre, suggesting one week is long enough time to observe the increased synapse formation in these cultures of from E18.5 neurons (Fig. 5d). Then, we pre-treated cultured hippocampal neurons with the function-blocking anti-Ryk monoclonal antibody generated by our lab for 2 hours before adding the Aβ oligomers ^35^. We found that Aβ oligomers failed reduce synapse numbers in the presence of the Ryk antibody (Fig. 5e,f). Again, during the 14 hours of time, Ryk antibody itself did not cause any change in synapse numbers in these cultures (Fig. 5f). Therefore, together with Vangl2, Ryk, activated by Wnt5a, is also required for Aβ oligomer-induced loss of glutamatergic synapses. As Aβ oligomers do not bind to Ryk (Extended data Fig. 9), we propose that Wnt/Ryk and Vangl2 work as one signaling axis which causes synapse disassembly and Aβ oligomers enhance their function.

**Fig. 5.**
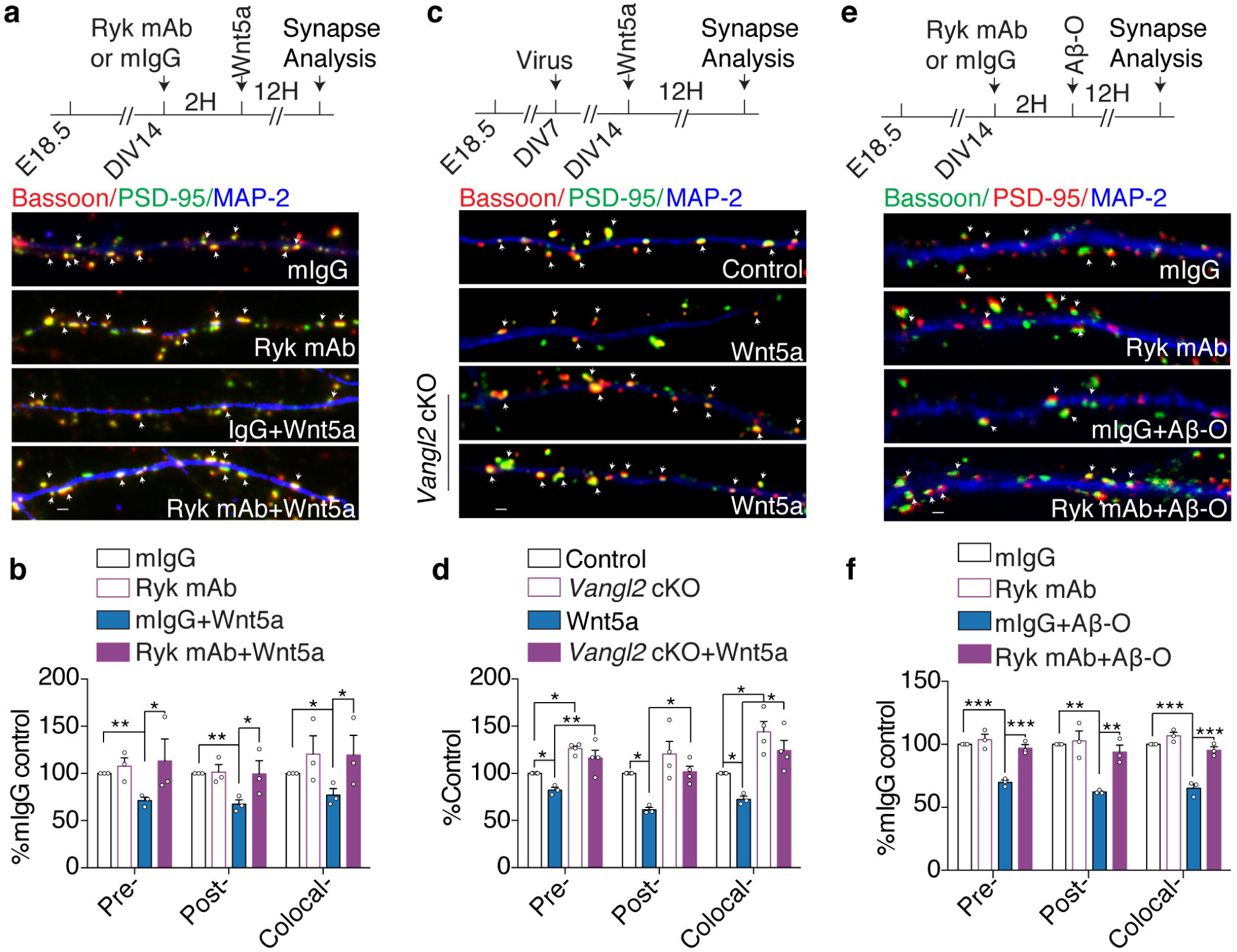
The Wnt/Ryk/Vangl2 signaling axis mediates synapse loss induced by oligomeric Aβ. **a-b**, Representative images and quantification of synaptic puncta (arrowheads) testing the effect of the Ryk antibody on Wnt5a-induced synapse reduction in cultured hippocampal neurons. n=3 experiments (n=27 neurons in IgG control, n=22 neurons in Ryk antibody, n=24 in IgG+Wnt5a, and n=20 neurons in Ryk antibody+Wnt5a). **c-d,** Representative images and quantification of synaptic puncta (arrowheads) testing the role of *Vangl2* in Wnt5a-induced synapse reduction in cultured hippocampal neurons. n=3 WT mice and n=4 *Vangl2 cKO* mice. **e-f,** Representative images and quantification of synaptic puncta (arrowheads) testing the effect if the Ryk antibody in oligomeric Aβ-induced synapse reduction in cultured hippocampal neurons. n=3 experiments (n=26 neurons in IgG control, n=33 neurons in Ryk antibody, n=34 neurons in oligomeric Aβ, and n=39 neurons in Ryk antibody+ oligomeric Aβ). \**P* < 0.05, \*\**P* < 0.01 and \*\*\**P* < 0.001. One-way ANOVA. Scale bar: 1 μm. Error bars represent SEM.

To ask whether the Wnt-Ryk signaling is required for Aβ oligomer-induced synapse loss in a mouse model of Alzheimer’s disease, we intracerebroventricularly infused the aforementioned function-blocking anti-Ryk monoclonal antibody into the 8-week-old *5XFAD* mouse (Fig. 6a) ^35^. Ryk antibody was infused via an osmotic minipump for 14 days and synapse numbers were analyzed in blind. The Ryk monoclonal antibody blocked the loss of glutamatergic synapses in *5XFAD* transgenic mice (Fig. 6b,c).

**Fig. 6.**
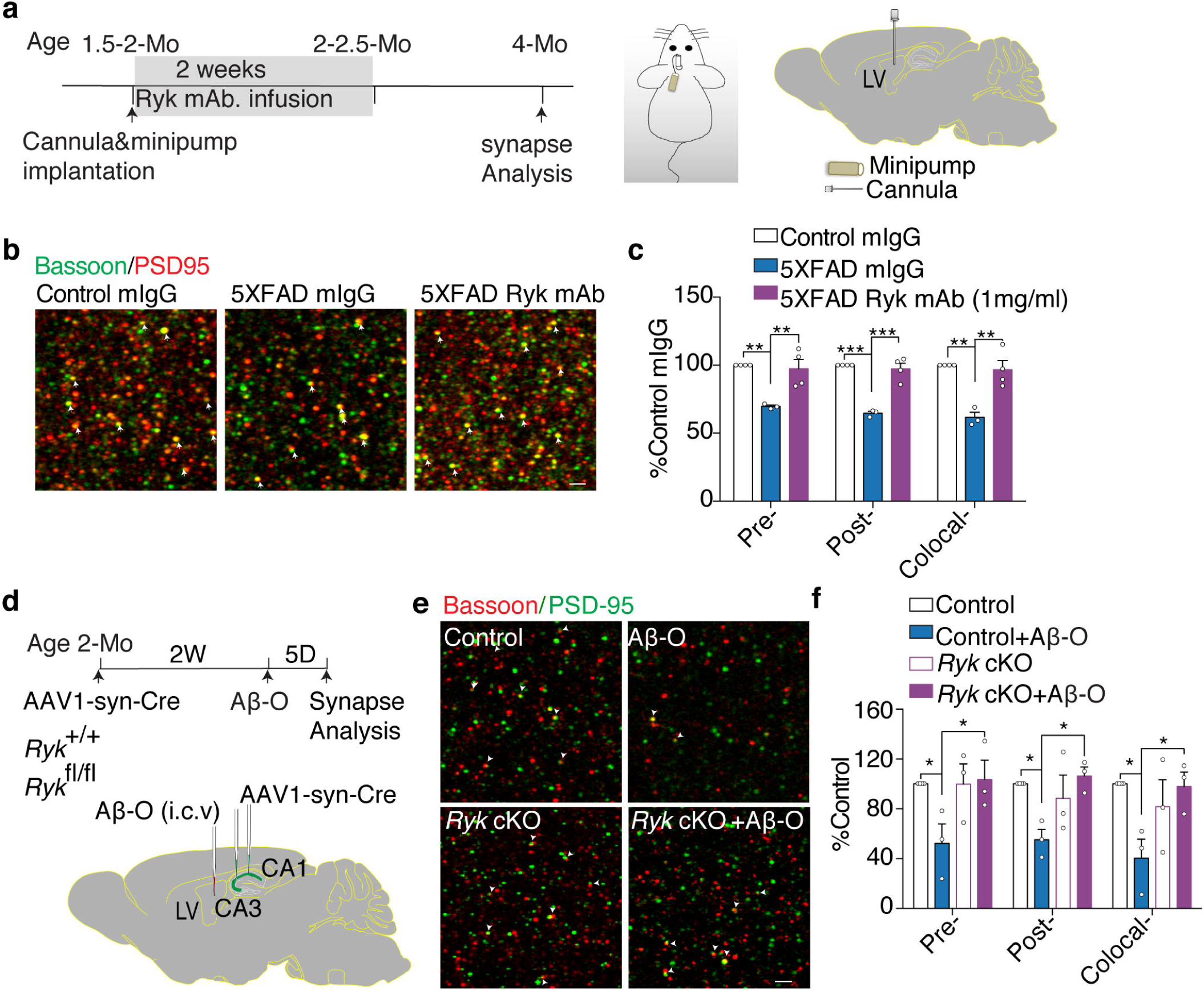
Ryk is required for oligomeric Aβ -mediated synapse loss *in vivo*. **a,** Timeline and schematic of intracerebroventricular infusion of the Ryk monoclonal antibody. Cannula and pre-infused osmotic minipumps were implanted at ∼2 months old. Minipumps were removed 2 weeks later. **b-c,** Staining and quantification of glutamatergic synapses in *5XFAD* mice infused with the Ryk monoclonal antibody. **d,** Schematics illustrating the experimental design. *AAV1-hSyn-eGFP-Cre* virus was injected into CA1 and CA3 regions of the hippocampus bilaterally. 2 weeks later, Aβ oligomers were injected into the lateral ventricular for 5 days. 5 days later, animals were fixed with perfusion and sectioning and stained with synaptic markers. **e-f,** Representative images and quantification of synaptic puncta detected by costaining for Bassoon (red)- and PSD95 (green)-immunoreactive (arrowheads) in stratum radiatum. n=4 for *Ryk*^+/+^ mice, n=3 for *Ryk*^+/+^ mice with oligomeric Aβ injection, n=3 for *Ryk cKO* mice and n=3 for *Ryk cKO* mice with oligomeric Aβ injection. \**P* < 0.05, One-way ANOVA. \*\*\**P* < 0.001. Scale bar: 1 μm in (**b**) and sale bar: 2 μm in (**e**). Error bars represent SEM.

To further confirm the role of *Ryk in vivo*, we injected of Aβ oligomers into the *Ryk cKO* mice previously generated in our lab and analyzed synapse numbers in blind ^35^. Because Ryk is likely involved on both the presynaptic and the postsynaptic side, *AAV1-hSyn-eGFP-Cre* was injected into the hippocampal CA1 and CA3 region of 8-week-old mice to conditionally knockout *Ryk* in adulthood. In *Ryk*^+/+^ (Control), Aβ oligomers induced a 60% reduction of synapse numbers. However, the synapse numbers were not reduced in the *Ryk cKO* mice with Aβ oligomers injected intracerebroventricularly (Fig. 6d-f).

### *Ryk cKO* prevents the loss of synapses and preserves cognitive function in *5XFAD* mice

To further characterize the role of *Ryk* on synapse loss in the *5XFAD* mice using a genetic approach, we crossed the *Ryk* cKO with *5xFAD*. *AAV1-hSyn-eGFP-Cre* was injected into the hippocampal CA1 and CA3 region of 8-week-old mice to conditionally knockout *Ryk* in adulthood. Glutamatergic synapse numbers were analyzed two months later (Fig. 7a). These experiments were done in double blind. We found that *Ryk cKO* can prevent the synapse loss (Fig. 7b,c). We analyzed the Ceslr3-positive synapse and found that the protected synapses are Celsr3-positive (Fig. 7d). Unlike in *Vangl2 cKO*, *Ryk cKO* itself showed a statistically insignificant (*p*=0.107) trend of increase of synapse numbers 2 months after the conditional knockout. Comparing with *5XFAD*, *Ryk cKO* lead to a 400% increase of synapse numbers in *5XFAD*, and preserved the synapse numbers to the control level (Fig. 7b,c). We noted that, compared to *Vangl2 cKO* (Fig. 1i,j), Ryk *cKO* rescued more synapses (Fig. 7b,c). This may be due to the differences of knockout efficiency of *Vangl2* and *Ryk* floxed alleles.

**Fig. 7.**
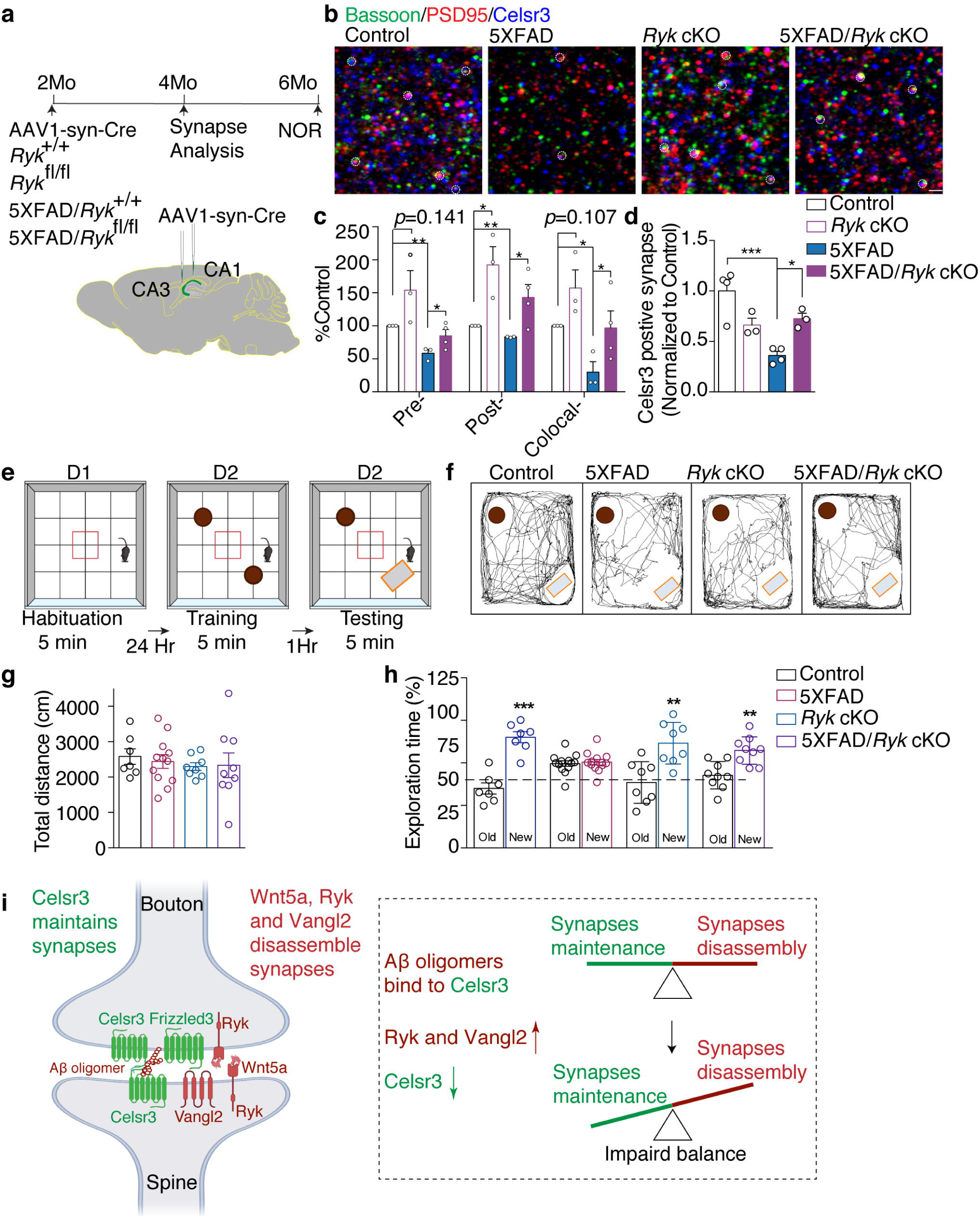
Function blocking Ryk antibody and *Ryk cKO* can prevent synapse loss and preserve cognitive function in *5XFAD* mice. **a**, Schematics for timeline and experimental design. *AAV1-hSyn-eGFP-Cre* virus was injected into CA1 and CA3 region of the hippocampus bilaterally. Animals were fixed with perfusion and sectioning and stained with synaptic markers at 4 months of age. A separate set of animals were injected and subjected to NOR at 6 months of age. **b-c,** Representative images and quantification of synapse numbers. **d,** Quantification of Celsr3-positive glutamatergic synapse (Celsr3 colocalized with PSD95 and bassoon). **e,** Schematic showing the design of novel object recognition (NOR). Mice were subjected to an open arena for three trails to evaluate the memory of objects. **f,** Trajectories of mice in the NOR test session. **g,** Quantification of locomotion. **h,** Quantification of NOR. Student *t*-test. **i,** Schematic diagram showing the balance of Wnt/PCP signaling in synapse maintenance and the binding site of oligomeric Aβ. \**P* < 0.05, \*\**P* < 0.01 and \*\*\**P* < 0.001. Scale bar: 1 μm in (**b**). Error bars represent SEM.

To test whether *Ryk* conditional knockout can preserve cognitive function, we performed the novel object recognition (NOR) test with a new set of mice at 6 months of age (Fig. 7a). These studies were also done in double blind. Novel object recognition tests were carried out in an open field arena measuring 0.4 × 0.4 × 0.45 m^3^ (Fig. 7e). The total distance was recorded during the 5-minute open field session as locomotor activity; no differences were found among the groups (Fig. 7f,g). For the NOR test, the animals were placed at the center of the arena in the presence of two identical (familiar) objects for a 5-minute training session. Exploratory behavior (amount of time exploring each object) was recorded by experienced researchers. One hour after training, animals were replaced into the arena for the testing session, in which one of the objects had been replaced by an unfamiliar (novel) object. The time spent exploring the familiar and novel objects was measured. We found that *Ryk cKO* partially preserved the cognitive function without affecting total distance of the locomotion (Fig. 7f-h). It is worth noting that the *Ryk cKO* itself did not cause any behavioral defects but it rescued the impaired novel object recognition behavior of the *5XFAD* mice. In contrast, we found that *Vangl2* cKO itself causes behavioral defects and failed to rescue cognitive function in the *5XFAD* Mice (Extended data Fig. 10). Based on this, Ryk may be a better therapeutic target for intervention than Vangl2. This difference may be caused by the fact that Vangl2 is a core PCP component and may play a more direct role on the composition of specific synaptic components important for the function of the synapses, whereas Ryk is not a core PCP component and may generally regulate the level of PCP signaling and synapse numbers. Therefore, Ryk may simply control the “quantity” of synapses, whereas Vangl2 may control both the “quantity” and the “quality”.

## DISCUSSION

PCP components are localized in the pre-and postsynaptic compartments in glutamatergic synapses and are the key regulators of glutamatergic synapse formation ^24^. We show here that PCP proteins are localized in adult synapses and continue to regulate synapse numbers in the mature nervous system and Celsr3 and the Wnt/PCP signaling pathway are a direct target of Aβ oligomers in causing synapse degeneration. Vangl2 disrupts the intercellular complex of Celsr3/Frizzled3-Celsr3 in neighboring cells. Aβ oligomers bind to three of the Celsr3 domains, one of which mediates the formation of Frizzled3/Celsr3 complex and thus disrupt this intercellular complex by weakening the interaction between Frizzled3 and Celsr3 on the presynaptic membrane. This function of PCP components in synapse assembly and disassembly is regulated by Wnt/Ryk signaling. *Ryk cKO* can protect synapses and preserve cognitive function in the *5XFAD* mice (Fig. 7i). Although Aβ oligomers themselves are not able to disrupt the complex, they cause synapse loss by tipping the balance of the PCP signaling components, allowing the Wnt/Ryk/Vangl2 signaling axis to more efficiently disassemble glutamatergic synapse and impair cognitive function.

Several receptors, including cellular prion protein (PrP^C^), EphB2 and paired immunoglobulin-like receptor B (PirB) or its human ortholog leukocyte immunoglobulin-like receptor B2 (LilrB2), have been reported for Aβ oligomers and regulate synaptic plasticity,. These receptors mediate the function of Aβ oligomers in altering synaptic function and plasticity but not synapse loss ^21 22 23^. Our study is the first to identify the binding protein of Aβ oligomers that directly mediates synapse loss. Our study also identifies a biological pathway, the Wnt/PCP pathway, that Aβ oligomers may target to negatively regulate neuronal activity by reducing the number of synapses acting as a negative feedback of neuronal activity. And this mechanism may become hijacked in pathological conditions, such as when Aβ is overproduced in patients of Alzheimer’s disease ^12^.

Aβ burden is generally considered a cause for both early onset and late onset Alzheimer’s disease. Our findings identified the Wnt/PCP pathway as a synaptic target of Aβ-associated synaptotoxicity, a process thought to start long before the appearance of cognitive symptoms and may provide new clues to understand pathogenesis. Our findings do not exclude the possibility that the PCP pathway may also be involved in tauopathy-mediated synapse degeneration, which likely occur at a later time, a topic we plan to explore in the future ^11^. Furthermore, our discovery of a direct synaptic target in mediating Aβ oligomer-induced synapse loss does not exclude or contradict the possibility of other mechanisms, such as by complement and microglia ^31^. However, the finding of the synaptic target of synaptotoxicity may provide new approaches for therapeutic design. Based on our current data, it appears that Ryk is a better therapeutic target than Vangl2 as *Vangl2 cKO* itself leads to behavioral deficits and cannot improve cognitive function and *Ryk cKO* itself does not lead to behavioral deficits and can significantly preserve cognitive function (Fig. 7h).

## METHODS

### Animals

All animal work in this research was approved by the University of California, San Diego (UCSD) Institutional Animal Care and Use Committee. The 5XFAD transgenic mice carrying the following five mutations: Swedish (K670N and M671L), Florida (I716V) and London (V717I) in human APP695 and human PS1 cDNA (M146L and L286V) under the transcriptional control of the neuron-specific Thy-1 promoter and were purchased from the Jackson Laboratory (Stock #34848) ^29^. 5XFAD mice were crossed with *Vangl2*^fl/fl^ (cKO), which was provided by Yingzi Yang at Harvard Medical School ^36^. CRISPR/Cas9 knockin mice were purchased from the Jackson Laboratory (Stock #024858).

### Aβ oligomer preparation

Human Aβ42 (AnaSpec) or human biotin-beta-Amyloid (1-42) (AnaSpec) was dissolved in dimethyl sulfoxide (DMSO), sonicated and diluted with F12 medium for Aβ monomerization to a concentration of 100 µM. For oligomerization, the solution was incubated for 24-26 hours at 4 °C, centrifuged at 16,000 x g for 20 min, and the supernatant was collected as oligomerized Aβ. The oligomerized Aβ42 preparations were analyzed via SDS-PAGE using 12% tris-glycine gels. 50 μg Aβ42 peptides were loaded and electrophoretically separated at 25 mA. Gels were transferred onto PVDF membrane. Forms of Aβ42 were detected using antibody 6E10 (BioLegend). 26.94±7.43% of Aβ42 existed as monomers (molecular weight 2.5-6.5 kD) (n=4); 2.06±0.41% as dimers (MW 6.5-11.5 kD); 17.84±0.97% as trimmers (MW 11.5-15.5 kD); 38.15±5.09% as tetramers (MW 15.5-20.5 kD) and 15.01±7.95% as high-n oligomers (Extended data Fig. 2).

### Intracerebroventricular (ICV) infusion of Aβ oligomer

Adult (2-3 months old) mice were deeply anaesthetized with an intraperitoneal injection of ketamine/xylazine cocktail until unresponsive to toe and tail pinch. Aβ oligomers (5 ng; volume 250nl) or PBS (volume 250 nl) was stereotaxically injected into bilateral ventricles (−0.1 mm anteroposterior, 1 mm mediolateral and -2.5 mm dorsoventral). 5 days after ICV injection, brains were harvested for immunohistochemistry.

### Intrahippocampal injection of AAV1-hSyn-eGFP-Cre

Adult Vangl2 cKO and littermates WT controls (2-3 months old) were deeply anaesthetized with an intraperitoneal injection of ketamine/xylazine cocktail until unresponsive to toe and tail pinch. AAV1-hSyn-eGFP-Cre (Addgene) was stereotaxically injected into bilateral hippocampal CA1 (160 nl per site).

### sgRNA design and expression analysis

The Brie database was used for the design of sgRNAs. To validate the efficiency of sgRNAs candidates, individual sgRNA was cloned into PX549-SpCas9 vector (Addgene #62988). Neuro2A cells were cultured on a 12-well plate. Cells were then transfected with 1μg sgRNA plasmid by using 1 mg/ml Polyethyleneimine MAX (Polyscience). Puromycin was used to selecte the transfected cells. Genomic DNA of Neuro2A was purified by using the cloroform-ethonal method. The amplification of target DNA fragment and efficiency testing of individual sgRNAs were performed with manufacture’s protocol of Surveyor assay.

Selected sgRNA sequences are as follows: *Celsr1* exon1: 5’- ACGTCTGGTGTGATCCGTAC-3’; *Celsr2* exon1: 5’-GTACACCGTTCGGCTCAACG-3’; Ceslr3 exon1 5’-CGTTCGGGTGTTATCAGCAC-3’. Three sgRNAs were cloned into all-in-one CRISPR/Cas9 vector using multiplex CRIPSR/Cas9 assembly system kit protocol. And then the multi-sgRNAs were cloned into the AAV vector (Addgene #60231) for virus package. qRT-PCR was used for virus titer measurement. The virus titer was ∼10^13^ GC/mL. AAV-sgRNA virus were stererotacically injected into the hippocampal CA1 region of the Cas9 mice.

### Hippocampal neuron culture

Hippocampi were dissected from E18.5 mice, and hippocampal neuron culture was performed as previously described ^24^. Briefly, cells ells were pelleted and resuspended in Neurobasal medium supplemented with 1% B27 (Invitrogen), penicillin/streptomycin (CellGro), and Gluta-MAX (Invitrogen) and plated on poly-D-lysine (Millipore) coated glass coverslips in a 24-well plate at a density of 2 Å∼ 10^4^ cells per square centimeter for immunostaining. Medium was changed every 4 days. Cultures were grown for 14 DIV at 37 °C with 5% carbon dioxide atmosphere.

### Immunofluorescence staining and image analysis of cultured neurons

For synaptic puncta density analysis in cultured hippocampal neurons, neurons on DIV14 were fixed for 20 min in 4% PFA. After fixation, cells were incubated in a blocking solution (1% bovine serum albumin, and 5% goat serum in Tris buffer saline solution (TBS) with 0.1% Triton X-100) for 1 h, and then stained overnight at 4 °C with primary antibodies chicken anti-MAP2 (neuronal marker; Abcam), guinea pig anti-Bassoon (presynaptic marker; Synaptic Systems) and goat anti-PSD-95 (postsynaptic marker; Millipore). After, cells were incubated with fluorochrome-conjugated secondary antibodies Alexa 488 anti-chicken, Alexa 647 anti-guinea pig and Alexa 568 anti-goat solution for 2 h at room temperature and mounted in mounting media. Z-stacked images were obtained with a Carl Zeiss microscope using a 63×oil-immersion objective. Three or more neurons with pyramidal morphology and at least two diameters’ distance from the neighboring neurons were selected per coverslip. Three coverslips were used for each group per experiment. Secondary dendrites were chosen for puncta analysis. Number of puncta was analyzed using the ImageJ Synapse Counter plug-in and the length of the dendrite was analyzed by ImageJ (NIH). Data in Fig. 1j, Fig. 2h, Fig. 6f, Fig. 7c were quantified in double blind. Data in Fig. 2d, Fig. 3i-l, Fig. 5b, Fig. 5d, Fig. 5f and were quantified in single blind.

### Immunofluorescence staining and image analysis of brain sections

For *in vivo* synaptic protein immunostaining, mice were deeply anesthetized with an intraperitoneal injection of ketamine/xylazine until unresponsive to toe and tail pinch and perfused with PBS followed by 4% PFA. Brains were removed and post-fixed in 4% PFA overnight at 4 °C. After, brains were cryoprotected in 30% sucrose for 2 days and coronal free-floating sections were prepared at 30 μm in a vibratome. The sections obtained were treated with 1% SDS for 5 min at room temperature for antigen retrieval, incubated in a blocking solution (1% bovine serum albumin, and 5% goat serum in Tris buffer saline solution (TBS) with 0.1% Triton X-100) for 1.5 h, and then stained overnight at 4 °C with primary antibodies guinea pig anti-Bassoon (presynaptic marker; Synaptic Systems) and goat anti-PSD-95 (postsynaptic marker; Millipore). After, sections were incubated with fluorochrome-conjugated secondary antibodies Alexa 647 anti-guinea pig and Alexa 568 anti-goat solution for 2 hours at room temperature, counterstained with DAPI and mounted in mounting media. The synapses formed between the Schaffer collaterals and the hippocampal CA1 pyramidal neuron apical dendrites spanning the mouse stratum radiatum were imaged. Fluorescent z-stack images were acquired with an LSM510 Zeiss confocal microscope using a 63× oil-immersion objective with 2X zoom-in. Number of puncta were analyzed using the ImageJ Synapse Counter plug-in.

### Plasmid, inhibitors and antibodies

Celsr3-Flag, Fzd3-HA, Vangl2-Myc and tdTomato expressing constructs were described previously ^37, 38^. Sulfo-NHS-LC-Biotin was purchased from Pierce. The antibodies used in this study include α-Vangl2 (Santa Cruz), α-Celsr3 (Rabbit polyclonal antibodies were generated by the Zou lab), α-Flag (Sigma), α- GAPDH (Chemicon), α-Insulin Rβ (Santa Cruz) and α-HA (Covance).

To generate truncated Celsr3 constructs, full-length Celsr3 extracellular domain is amplified by PCR, digested with EcoRV/NheI, and subcloned into pCAGEN vector using primers as follows:

ΔEGF/Lam_Celsr3 Forward primer 1: 5’-GATCGATATC TTCTCTGGAGAGCTCACAGC-3’

ΔEGF/Lam_Celsr3 Reverse primer 1: 5’-GCAGGCATCGTA AAAGGGCAGCACGTCGAG-3’

ΔEGF/Lam_Celsr3 Forward primer 2: 5’-GTGCTGCCCTTT TACGATGCCTGCCCCAAG-3’

ΔEGF/Lam_Celsr3 Forward primer: 5’-GATCGCTAGCAAGTAGGCCAGCAAG-3’

ΔEGF1_Celsr3 Forward primer: 5’- TGCTGCCCTTTACAGAGCTCGACCTCTGTTAC-3’

ΔEGF1_Celsr3 Reverse primer: 5’-CGAGCTCTGTAAAGGGCAGCACGTCGAG-3’

ΔEGF2_Celsr3 Forward primer: 5’- TCTGTGAGACACTGGACACTGAAGCTGGACG-3’

ΔEGF2_Celsr3 Reverse primer: 5’- TCAGTGTCCAGTGTCTCACAAGAAGTCTCCCG-3’

ΔEGF3_Celsr3 Forward primer: 5’-GCTGGACACTGTGGCCGCACGCTCCTTTC-3’

ΔEGF3_Celsr3 Reverse primer: 5’-GTGCGGCCACAGTGTCCAGCTCGCAGTC-3’

ΔLaminin G1_Celsr3 Forward primer: 5’- ACGCTGTGAGCAGGCCAAGTCACACTTTTGTG-3’

ΔLaminin G1_Celsr3 Reverse primer: 5’- ACTTGGCCTGCTCACAGCGTGGACCATC-3’

ΔEGF4_Celsr3 Forward primer: 5’-AGGCTGCCAGCTCACAATGGCCCATCCCTAC- 3’

ΔEGF4_Celsr3 Reverse primer: CCATTGTGAGCTGGCAGCCTGCCATAGTG-3’

ΔLaminin G2_Celsr3 Forward primer: 5’- CTGTCGACTCACTGTGACCAACCCCTGTG-3’

ΔLaminin G2_Celsr3 Reverse primer: 5’- TGGTCACAGTGAGTCGACAGTCTTTGCCACC-3’

ΔEGF5_Celsr3 Forward primer: 5’-TGGCTGTACTGATGCCTGCCTCCTGAACC-3’

ΔEGF5_Celsr3 Reverse primer: 5’- GGCAGGCATCAGTACAGCCAGGCTCCACATTC-3’

ΔEGF6_Celsr3 Forward primer: 5’- AGGCTGTGTGTATTTTGGTCAGCACTGTGAGCAC-3’

ΔEGF6_Celsr3 Reverse primer: 5’- GCTGACCAAAATACACACAGCCTGGGCCATAG-3’

ΔEGF7_Celsr3 Forward primer: 5’-TGTGAGTGGCAAGACGAATGGCCAGTGCC-3’

ΔEGF7_Celsr3 Reverse primer: 5’-CCATTCGTCTTGCCACTCACAGTCACAAG-3’

ΔEGF8_Celsr3 Forward primer: 5-CAACTGCAACCCCCACAGCGGGCAGTG-3’

ΔEGF8_Celsr3 Reverse primer: 5’-CTGTGGGGGTTGCAGTTGGGGTCAAAGC-3’

ΔLaminin EGF_Celsr3 Reverse primer: 5’- GCATCGTAGAGTGGGAGGCATGAGTCACTG-3’

ΔLaminin EGF_Celsr3 Forward primer: 5’- ATGCCACCCACTCTACGATGCCTGCCCCAAG-3’

### HEK293T cell transfection

HEK293T cells were purchased from ATCC and maintained in Dulbecco’s modified Eagle’s medium (DMEM) containing 10% fetal bovine serum. Transfection of HEK293T cells was carried out using 1 mg/ml Polyethyleneimine MAX (Polyscience). Mycoplasma contamination was monitored by DAPI staining.

### Aβ oligomer Binding assay

HEK293 cells were transiently transfected (polyethylenimine) with expression vectors encoding TdTomato, Celsr3-Flag or control empty vectors (pCAGEN). Two days post-transfection, cells were treated with biotinylated Aβ oligomer for 2 h at 37°C, washed twice, and fixed with 4% PFA for 20 min, blocked with 5% donkey serum in PBS with 0.1% Triton X-100. The bound Aβ peptides were visualized with streptavidin-Alexa fluorophore conjugates (Alexa 488). DAPI was used to counterstain cell nuclei; TdTomato was used to monitor construct transfection. Anti-flag antibody was used to stain Celsr3. Fluorescent images were captured with Zeiss LSM 880 fast Airyscan using a 63× oil-immersion objective. For saturation binding assays and calculation of the dissociation constant, K*_d_*, 48 h after transfection, cells were incubated with serial dilutions of biotinylated Aβ oligomer for 2 h at 37 °C. Control experiments were performed using the maximum amount of biotinylated Aβ oligomer but HEK293T cells were transiently transfected with the pCAGEN vector. Cells were washed and fixed with 4% PFA for 10 min, washed with 0.1% Triton X-100/PBS and incubated for 30 min with extra-avidin peroxidase (Sigma-Aldrich). Cells were then extensively washed and bound peroxidase was quantified using TMB substrate (Thermo Fisher Scientific). The reaction was terminated by addition of TMB stop solution (Thermo Fisher Scientific). Absorbance was read at 450 nm in a UV-Vis microplate reader. Non-specific binding was determined in the presence of 100 μm non-biotinylated Aβ oligomer and specific binding was calculated by subtracting absorbance values for nonspecific binding from total binding values.

### Surface biotinylation assay to characterize cell surface expression levels of Celsr3 and Celsr3 deletion constructs (Extended data Fig. 4)

The surface biotinylation and NeutrAvidin pull down methods have been described previously ^37, 38^. Briefly, 48 hr after transfection with indicated plasmids, HEK293T cells (seeded on 20 μg/ml PDL-coated six-well plate) cells were washed with ice-cold PBS (pH 8.0) three times and incubated with 1 mg/ml Sulfo-NHS-LC-Biotin (ThermoFisher Scientific)/PBS for 2 min at room temperature to initiate the reaction, followed by incubation on ice for 1 hr. After quenching active biotin by washing with ice-cold 100 mM Glycine/PBS twice followed by normal ice-cold PBS, the cell lysates were incubated with NeutrAvidin agarose for 2 hr and then precipitated. For quantification, three independent experiments were performed, and the band intensity was quantified with ImageJ (NIH).

### Coimmunoprecipitation

48 hr after transfection with the indicated plasmids, HEK293T cells were lysed with IP buffer (20 mM Tris HCl (pH 7.5), 150 mM NaCl, 1 mM EGTA, 5 mM NaF, 10 mM β- glycerophosphate, 1 mM Na_3_VO_4_, 1 mM DTT and protease inhibitor cocktail (SIGMA), 0.1% TX-100). Lysates were immunoprecipitated with anti-HA, anti-Myc or anti-Flag antibodies and with protein A/G agarose (Santa Cruz). Experiments were repeated three times and showed similar results.

### Ryk monoclonal antibody infusion

Osmotic minipump (Model 1002, Alzet, Cupertino, CA) were pre-filled with either 100 µl mouse IgG or Ryk monoclonal IgG (clone 25.5.5 generated against the ectodomain of Ryk, amino acid range 90-183, by Antibody Solutions, Santa Clara, CA) in artificial cerebrospinal fluid. A subcutaneously implanted osmotic minipump connected by polyvinylchloride tubing to a stainless-steel cannula stereotaxically implanted into the lateral ventricle (Brain Infusion Kit 1; Alzet). Mice were randomly selected for mouse IgG or Ryk monoclonal IgG treatment. Osmotic minipumps were removed after 14 days.

### Glutathione S-transferase fusion protein generation and biotinylated Aβ-pull down assay

Glutathione S-transferase (GST) or GST fusion of the EGF7 domain and EGF8 domain of Celsr3 (GST-EGF7 and GST-EGF8) were generated using pGEX4T-1. All GST fusions were expressed in BL21 *Escherichia coli* and purified with glutathione-Sepharose 4B (GSH beads; GE Healthcare). 100µM of biotinylated- Aβ oligomer and 200µg of GST, GST-EGF7, and GST-EGF8 were incubated with NeutrAvidin agarose slurry for 2.5 hours in cold room and then precipitated. The protein before precipitated and after precipitated were analyzed by Western blotting.

### Novel objective recognition behavioral testing

Object recognition tests were carried out in double blind in an open field arena measuring 0.4 × 0.4 × 0.45 m^3^. The total distance was recorded during a 5-minute open field session as locomotor activity; no differences were found among the groups. For NOR test, the animals were placed at the center of the arena in the presence of two identical (familiar) objects for a 5-minute training session trained. Exploratory behavior (amount of time exploring each object) was recorded by experienced researchers. One hour after training, animals were replaced into the arena for the testing session, in which one of the objects had been replaced by an unfamiliar (novel) object. The time spent exploring familiar and novel objects was measured using SMART video tracking software (PanLab Harvard Apparatus, Holliston, MA, USA). Animals that had a total exploration time below 8 s were excluded from the NOR tests. Results are expressed as the percentage of time exploring each object during the training or testing session, and were analyzed using a one-sample Student’s *t*- test comparing the mean exploration time for each object with the fixed value of 50% (chance level).

### Statistical analyses

Comparisons between multiple experimental groups were performed by one-way ANOVA followed by Tukey-Kramer post-hoc test, when appropriate. Comparisons between two experimental groups were performed by Student’s *t* test. All statistical analyses were performed using GraphPad Prism software (La Jolla, California, USA). A value of *p*<0.05 was considered significant.

## ACKNOWLEDGMENTS

We would like to thank Kuanhong Wang, Zhigang He and members of the Zou lab for critical reading and comments on the manuscript. This project was supported by <u>RO1 MH116667 to Y.Z.</u> Airyscan confocal microscopy imaging was performed at UCSD School of Medicine Light Microscopy Facility (Grant P30 NS047101).

## AUTHOR CONTRIBUTIONS

Y.Z. and B.F. designed the experiments; B.F., A.E.F., R.Y.T., A.S.G., J.W., and Y.R.L. performed all experiments under the supervision of Y.Z; B.F., R.Y.T., A.E.F., and Y.Z. analyzed and interpreted data; B.F. and Y.Z. wrote the paper; and all authors read and commented on the manuscript.

## COMPETING INTERESTS

Yimin Zou is the founder of VersaPeutics. The terms of this arrangement have been reviewed and approved by the University of California, San Diego in accordance with its conflict of interest policies.

**Extended data Fig. 1 Celsr is required for synapse maintenance in adulthood, Related to Fig. 1.**

**a,** Sequence of selected sgRNAs and gel images of PCR products amplified from the target sites of Celsr1, Celsr2 and Celsr3 in Neuro2A cells transfected by SpCas9 and corresponding sgRNA. **b,** Western blotting showing the deletion efficiency of the selected sgRNAs in cultured hippocampal neurons. **c-d,** Representative images and quantification of synaptic puncta detected by costaining for Bassoon (red)- and PSD95 (green)- immunoreactive in stratum radiatum. \*\**P* < 0.01 and \*\*\**P* < 0.001. Student *t*-test. Scale bar: 2 μm in (C). Error bars represent SEM.

**Extended data Fig. 2 Characterize of Aβ oligomers, Related to Fig. 2.**

Total Aβ42 oligomers were separated from Aβ42 monomer in 12% SDS-PAGE Gel. Aβ42 oligomers were composed by different sizes of oligomers ranging from 2-mer to 4-mer.

**Extended data Fig. 3 Aβ42 monomers doesn’t bind to Celsr3, Related to Fig. 3.**

Celsr3-Flag (red)-transfected HEK293T cells were incubated with monomeric Aβ42 (200 nM total peptide), and monomeric Aβ42 was labeled green. Scale bar 10 μm.

**Extended data Fig. 4 Surface expression of truncated Celsr3. Related to Fig. 3. a,** Surface expression of ΔEGF/Lam_Celsr3 and Celsr3. Cell surface proteins were labeled with biotin and then precipitated with Neutravidin agarose. Precipitants and total lysates were subject to immunoblotting with the indicated antibodies. **b,** Surface expression of Celsr3 and with individual domain deletion. **c,** Truncated Celsr3-Flag (red) transfected HEK293T cells were treated with oligomeric Aβ42 (200 nM total peptide, monomer equivalent), and bound oligomeric Aβ42 (green) was visualized using 488-conjugated streptavidin. Scale bar 10 μm.

**Extended data Fig. 5 Aβ oligomers binds to human-Celsr3, Related to Fig. 3.**

**a,** Amino acid alignment of Laminin G1, EGF 7 and EGF 8 domains of *h*Celsr3 and *m*Celsr3. **b,** Binding of oligomeric Aβ42 (200 nM total peptide, monomer equivalent) with *h*Celsr3-Flag-transfected or truncated *h*Celsr3-Flag-tramsfected HEK203T cells. Bound oligomeric Aβ42 (green) was visualized using 488-conjugated streptavidin. Scale bar 10 μm.

**Extended data Fig. 6 Interaction between Vangl2 and Celsr3 is not affected by the oligomeric Aβ42 binding domain, Related to Fig. 3.**

**a,** Co-IP assays testing the interaction between Vangl2 and Celsr3 or with truncated Celsr3. **b,** Quantification data of the expression level of co-IPed Vangl2. \**P* < 0.05. One-way ANOVA. One-way ANOVA. Means ± SEM.

**Extended data Fig. 7 Celsr3 in the neighboring cell does not affect the interaction between Frizzled3 and Celsr3 in the same cell, Related to Fig. 4.**

**a,** Co-IP assays testing the interaction between Celsr3 and Frizzled3 with Celsr3 in the neighboring cell. **b,** Quantification data of the expression level of co-IPed Celsr3. Student’s *t* test.

**Extended data Fig. 8 Oligomeric Aβ42 on the interaction of Frizzled3 and Celsr3, Related to Fig. 4.**

**a-b,** IP assays testing the interaction between Celsr3 and Frizzled3 transfected in the same cell shows no significant difference with or without oligomeric Aβ42. **d-f,** Oligomeric Aβ42 and Vangl2 on the interaction between Frizzled3 and Celsr3 on the same. **g-i,** Oligomeric Aβ42 on interaction between Frizzled3 in one cell and Celsr3 on the other neighboring cell. \**P* < 0.05 and \*\*\**P* < 0.001. One-way ANOVA. Western blot results are representative of three (**c**) and four (**f** and **i**) biological replicates. Error bars represent SEM.

**Extended data Fig. 9 Oligomeric Aβ42 does not bind to Ryk, Related to Fig. 5.**

HEK293T cells transfected with Ryk-HA or control vector (pCAGEN) were incubated with oligomeric Aβ42 (200 nM total peptide, monomer equivalent). Bound oligomeric Aβ42 (green) was visualized using 488-conjugated streptavidin.

**Extended data Fig. 10 *Vangl2 cKO* caused defects in novel object recognition and did not rescue the phenotype of *5XFAD*, Related to Fig. 7.**

**a,** Quantification of locomotion. **b,** Quantification of NOR testing. \**P* < 0.05 and \*\*\**P* < 0.001. Student *t*-test. Error bars represent SEM.

